# Multielectrode array characterization of human induced pluripotent stem cell derived neurons in co-culture with primary human astrocytes

**DOI:** 10.1101/2024.03.04.583341

**Authors:** Maddie R. Lemieux, Bernhard Freigassner, Zahra Thathey, Mark R. Opp, Charles A Hoeffer, Christopher D. Link

**Author notes:** Corresponding authors: Maddie Lemieux (ML) Chris Link (CL).

## Abstract

Human induced pluripotent stem cells (hiPSCs) derived into neurons offer a powerful *in vitro* model to study cellular processes. One method to characterize functional network properties of these cells is using multielectrode arrays (MEAs). MEAs can measure the electrophysiological activity of cellular cultures for extended periods of time without disruption. Here we used WTC11 hiPSCs with a doxycycline-inducible neurogenin 2 (NGN2) transgene differentiated into neurons co-cultured with primary human astrocytes. We achieved a synchrony index ∼0.9 in as little as six-weeks with a mean firing rate of ∼13 Hz. Previous reports show that derived 3D brain organoids can take several months to achieve similar strong network burst synchrony. We also used this co-culture to model aspects of sporadic Alzheimer’s disease by mimicking blood-brain barrier breakdown using a human serum. Our fully human co-culture achieved strong network burst synchrony in a fraction of the time of previous reports, making it an excellent first pass, high-throughput method for studying network properties and neurodegenerative diseases.

## Introduction

The brain can respond to different stimuli in numerous ways depending on cell specialization and connectivity of neuronal circuits [1]. Studying these response patterns is important for understanding normal brain function and neurodegenerative disease states. Multielectrode arrays (MEAs) are a powerful tool that can be used to measure network electrophysiological changes. A common use of MEAs is to measure ex-vivo brain slice plasticity. However, a less exploited benefit of this system is the ability to characterize *in vitro* neuronal networks without cellular disruption [2,3]. Using MEAs one can record extracellular action potentials and local field potentials of populations of neurons for long periods of time [4]. These readouts can advance the neuroscience field due to their sensitivity and ability to mimic aspects of clinical data [5].

One MEA approach is to use human neurons derived from induced pluripotent stem cells (iPSCs). This model has advantages compared to using rodent cells as it provides phenotypic relevance to human brain neural networks and eliminates the species gap [6]. Human neuronal cultures begin with asynchronous, sporadic spiking due to lack of network interconnectivity [3]. The firing of one neuron may occur as a single spike or in clusters called bursts. Bursts are described as a period of intense spiking activity followed by a period of quiescence, and a network burst is the coordination of these spiking patterns across multiple cells [7]. As the culture develops, a synchronous network forms, suggesting the integration of individual neurons into an organized circuit [4]. Some critical parameters measured by multielectrode arrays include mean firing rate, network burst frequency, synchrony, and the coefficient of variation for the network interburst interval (IBI). The synchronization of cultures refers to the coordinated or simultaneous spiking between cells, and the coefficient of variation for the network IBI is a measure of network burst regularity. To achieve this strong, synchronous network bursting pattern, iPSCs derived from two-dimensional neuron monocultures need to be cultured with supporting cells such as astrocytes [3,8].

One major limiting factor of human based *in vitro* MEA studies is the time it takes to achieve network synchrony. Rodent cell models acquire this property within a few weeks while human cultures can take several months [3, 9]. Specifically, it has been found that human organoids take more than 30 weeks to reach a mean firing rate above 10 Hz and a synchrony index of 0.8 [10]. Two-dimensional cultures using human iPSC neurogenin 2 (NGN2) inducible neurons only achieved this firing rate when cultured with rodent astrocytes and not with human iPSC-derived astrocytes [11,12].

MEAs have also been used to model neurodegenerative disease states. Modeling sporadic Alzheimer’s disease (AD) is of significant relevance as it is the most common form of neurodegenerative disease [13]. Most *in vivo* animal approaches, however, use transgenic rodents, which only capture the genetic aspect of onset as amyloid beta plaques or tau tangles do not spontaneously form in aged rodents [13]. Recently, an *in vitro* method has been established by Chen et al. [14]. They treated human organoids with 10% human serum to replicate the breakdown of the blood-brain barrier seen in Alzheimer’s disease patients. After treatment, they observed AD-like pathologies, including increased amyloid beta aggregates and phosphorylated microtubule-associated tau protein levels. They also noted synaptic loss and impaired neural networks observed by MEA analysis, including reduced mean firing, synchrony, and bursts. In addition, single-cell transcriptomic analysis revealed immune activation of astrocytes after relatively chronic exposure [14]. This model provides a basis for future pharmacological interventions to be tested in a high throughput manner. However, as discussed previously, a limiting factor of this human organoid *in vitro* system is the amount of time it takes to achieve maturity of the culture.

Here, we present results of studies using fully humanized *in vitro* neuronal networks cultured on multielectrode arrays that were able to reproducibly achieve strong network burst synchrony in as little as six weeks. We used WTC11 human iPSCs containing a doxycycline-inducible neurogenin 2 (NGN2) transgene derived into neurons co-cultured with primary human astrocytes. The WTC11 cell line is particularly well characterized and is the cell line used for reporter iPSCs by the Allen Institute [15]. The NGN2-inducible derivative of this line can be quickly and reproducibly induced to make a population of largely excitatory neurons [16], which, for example, have been used both to model neurodegenerative disease [17] and as the basis for CRISPR-based screens [18]. In addition, we explored electrophysiological changes potentially associated with sporadic Alzheimer’s disease by treating our cultures with 10% human serum. Overall, we have established a fully human *in vitro* model that accelerates maturation time compared to existing methods that can be used to investigate neuronal network function in a high throughput manner.

## Materials and methods

### Maintenance and expansion of hiPSCS

WTC11 human induced pluripotent stem cells (iPSCs) containing a doxycycline inducible NGN2 transgene were cultured on 6-well Matrigel-coated plates as previously described by Fernandopulle et al. [16]. mTeSR plus media was changed daily or every other day until cells reached near 90% confluency. Cells were washed with PBS and split with 0.5 mM EDTA (room temperature for 5-7 minutes) for expansion or accutase (37⁰ C for 4 minutes) for single-cell differentiation.

### Generation of hiPSC derived neurons

hiPSCS were single cell split with accutase and plated on 10 cm Matrigel coated plates. The induction media consisted of DMEM/F12 with HEPES, N2 (1X, Gibco), NEAA (1X, Gibco), Glutamax (1X, Gibco), doxycycline (2 µg/mL), and ROCK inhibitor Y-27632 (10 µM). Media was changed daily for 3 days with only the initial plating media containing ROCK inhibitor. Cells were then detached with accutase, spun down at 250 x g for 5 minutes, resuspended, and either frozen or re-plated to complete maturation. Cortical maturation media contained BrainPhys neuronal medium (STEMCELL Technologies), B27 (1X, Gibco), BDNF (10 ng/mL, PeproTech), NT-3 (10 ng/mL, PeproTech), laminin (1µg/mL, Gibco), and doxycycline (2 µg/mL). Maturation was either completed on 8-well Permanox treated chamber slides or directly on multielectrode arrays. The minimum time for cells to reach maturity was 14 days.

### Astrocyte Culturing

Fetal human astrocytes (Lonza) were cultured in Astrocyte Media (ScienCell) on 6-well plates until 80-90% confluent. PBS was used to wash the cells before collection. Cells were detached using TrypLE express (Gibco) at 37⁰ C for 3-5 minutes. Following incubation, astrocyte media was added to stop the reaction. The cells were collected and centrifuged at 200- 300 x g for 5 minutes. Fresh astrocyte media was used to resuspend the cell pellet.

### Co-culture on multielectrode arrays

Multielectrode array 24-well plates containing 16 electrodes per well were obtained from Axion Biosystems (CytoView MEA plate). The electrodes were arranged in a 4x4 grid 350 µm apart, with each electrode diameter being 50 µm. The electrode recording area was 1.1 mm x 1.1 mm. Plates were sterile upon arrival and prepared for cell adhesion by treating the electrode field with 0.1 mg/mL Poly-D-Lysine at 37⁰ C for 4.5 hours. The wells were washed with water three times and dried in a biosafety cabinet overnight. 20 µg/mL laminin droplets were added to each field and incubated at 37⁰ C for at least one hour. A master mix of neural progenitor cells and astrocytes was made (90,000 NPCs: 45,000 astrocytes per well) and plated in 12 µL drops on each electrode field. The cell culture droplets were incubated at 37⁰ C for one hour, followed by adding 1 mL cortical media to fill the well. Water was added to the surrounding reservoirs to prevent media evaporation. Media changes occurred every 2-4 days by performing half media changes. Cells matured for about one month.

### Electrophysiology recordings

Spontaneous extracellular activity was acquired using Axion Biosystem Maestro Edge MEA. Under this instrumentation, cells were kept at 37⁰ C with 5% CO_2_ for the duration of the recordings. The Axion default sampling rate was 12.5 kHz/channel, and the signal was passed through the spontaneous filter (200 Hz-3 kHz). Recordings were taken for 6-minute durations. Analysis was done using the Axis plotting tool and neural metrics tool.

Unless otherwise specified, the spike threshold was set to ≥ 6 standard deviations above noise levels with active electrodes having at least 5 spikes/min for spontaneous recordings. Single electrode bursts had a minimum of 5 spikes with a max interspike interval of 100 milliseconds (ms). Network burst parameters were set to a minimum of 50 spikes per network burst with a max interspike interval of 10 ms. At least 35% of electrodes were coordinated to be considered a well with network bursts.

### Immunofluorescent staining

Cultures were grown in 8-well Permanox chamber slides and fixed at various time points between 14-33 days. Media was removed, and cells were fixed with 4% paraformaldehyde for 10 minutes. After fixation, cells were permeabilized with 0.1% Triton-X100 PBS solution for 5 minutes. The cultures were blocked with 5% bovine serum albumin (BSA) for 30 minutes, and primaries were then added and kept overnight at 4⁰ C. Secondary antibodies were added for 1 h at room temperature followed by DAPI (1 µg/mL) staining. Primary antibodies used were MAP2 (1:200, Millipore), β3-Tubulin (1:200, Cell Signaling Technology), and GFAP (1:500, Novus Biologics). Secondaries used were Alexa fluor 555 and 488, Invitrogen at 1: 500 or 1:1000 dilutions. Imaging was performed with Nikon A1R Laser Scanning Confocal.

### Statistical analysis

Data was collected and analyzed via the Axis plotting tool and neural metrics tool provided in the Axion Biosystem Maestro Edge software package. Statistical analyses were performed from the exported data files. Most data were analyzed using repeated measures ANOVA with an alpha value of 0.05 followed by Bonferonni corrections. Two-plate comparisons were made using independent t-tests with an alpha value of 0.05.

## Results

### Generation and characterization of human neuronal cultures

WTC11 human induced pluripotent stem cells (hiPSCs) containing a neurogenin 2 (NGN2) transgene were differentiated into neural progenitor cells (NPCs) using a 3-day, 2 µg/mL doxycycline induction as previously described [16]. Pre-differentiated NPCs were further matured into neurons using cortical maturation media (CM) for a minimum of 14 days. Cells were fixed and characterized with mature neuronal markers such as MAP2 and β3-Tubulin (Fig 1). In order to achieve optimal electrophysiological performance, NPCs were co-cultured with primary human astrocytes during maturation at a 2:1 ratio. These co-cultures were characterized with a neuronal MAP2 marker and an astrocyte marker GFAP.

**Figure 1.**
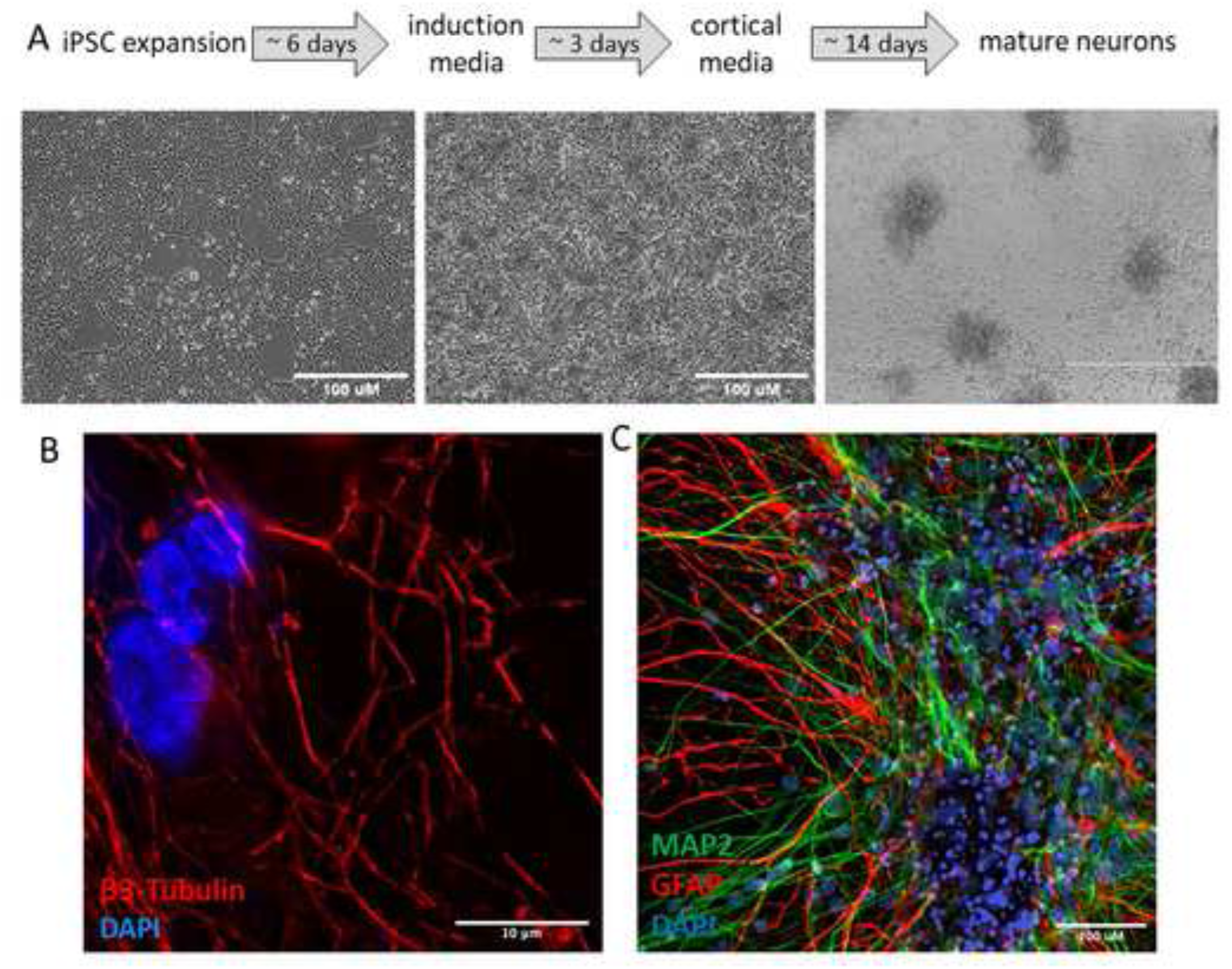
Generation and characterization of hiPSC derived neuron and astrocyte co-culture. (A) Workflow and timeline to produce mature neurons from WTC11 neurogenin 2 inducible hiPSCs (B) Mature neurons stained for β3-Tubulin (red) and DAPI (blue) (C) Co-culture stained for mature neuronal marker MAP2 (green), astrocyte marker (red) and DAPI (blue)

### Co-culture electrophysiological maturation

Multielectrode arrays were used to characterize the electrophysiological properties of cellular network formation. Co-cultures containing 90,000 NPCs and 45,000 astrocytes were plated on Axion Biosystem CytoView MEA 24-well plates. The cells were recorded and monitored visually for about 30 days before treatments. Recordings were taken daily starting on day 11 of maturation, and all 24-wells were averaged and quantified for specific parameters (Fig 2). Spontaneous firing was observed 11 days *in vitro* (DIV); however, there was no network activity. After 17 DIV, the mean firing rate increased more than 10-fold and almost all electrodes were active. A cellular network formed as seen by the synchronized network bursts appearing at a frequency of 0.11 Hz. The network interburst interval coefficient of variation was 0.44, and the synchrony index was 0.58 17 DIV. From this initiation of network activity, the co-culture continued to mature and stabilize until 33 DIV. This maturation is summarized in Fig 2. The average mean firing rate 26 DIV was 17.4 Hz with a network burst frequency of 0.16 Hz. The synchrony index was 0.82, and the network IBI coefficient of variation was 0.35. While the mean firing rate and network burst frequency peaked at this timepoint, the synchrony index and network IBI coefficient of variation continued to improve. At 33 days *in vitro*, the cultures were considered mature with an average mean firing rate of 12.9 Hz and synchrony index of 0.91. The average network burst frequency was 0.06 Hz with the network burst duration reaching its peak of 1.03 seconds. In addition, the cultures were considered stable with a network IBI coefficient of variation of 0.23, indicating the regularity of network bursting.

**Figure 2.**
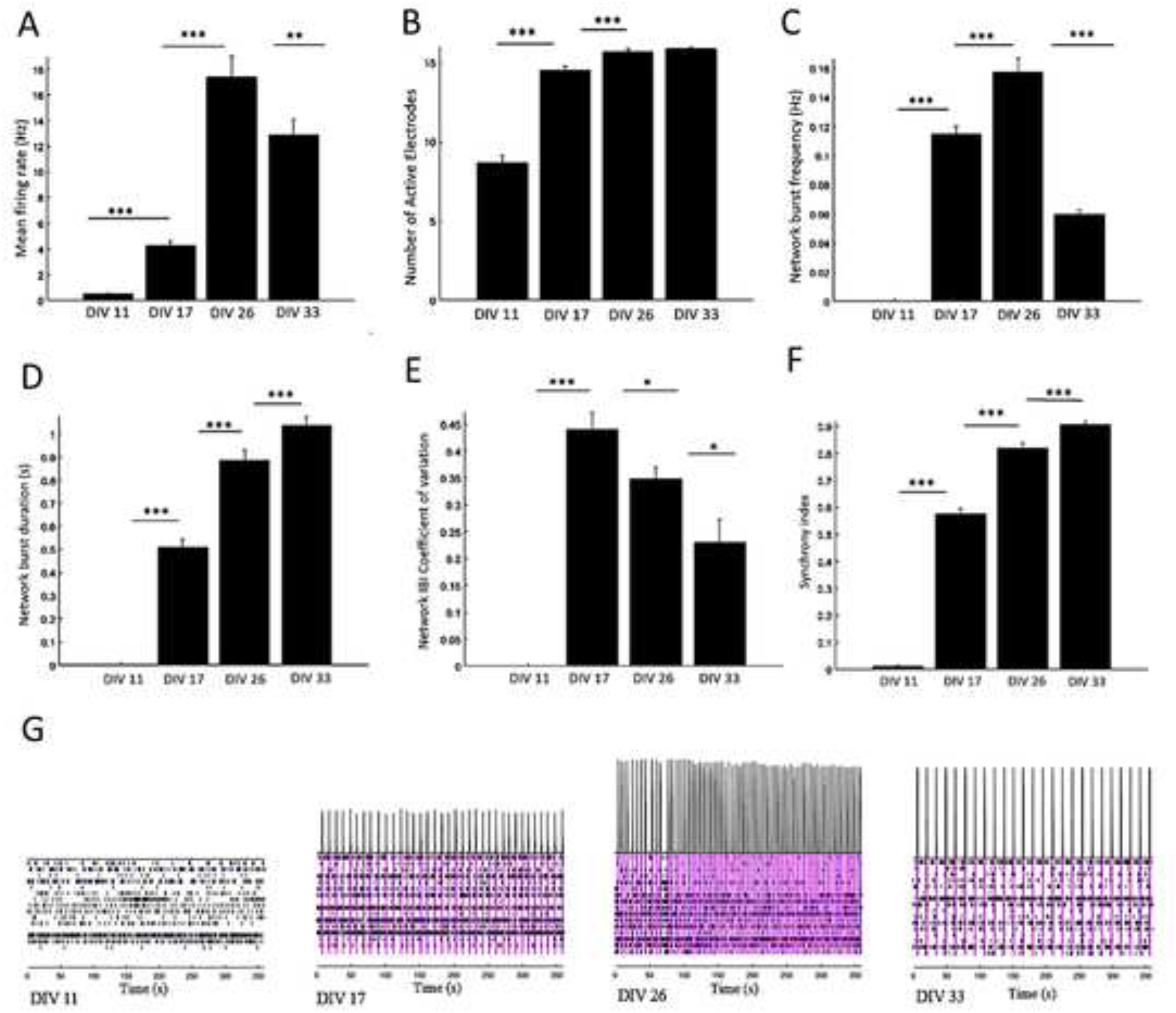
Co-culture electrophysiological maturation. 24-well quantification of (A) Mean firing rate (Hz), (B) Number of active electrodes, (C) Network burst frequency (Hz), (D) Network burst duration (s), (E) Network interburst interval coefficient of variation, and (F) Synchrony index on DIV 11, 17, 26, and 33. (G) Representative raster plots. Statistical analysis was performed with repeated measures ANOVA and Bonferroni corrections Error bar: SEM. N=24 independent wells. p-value < 0.05 *, p-value < 0.01 **, p-value < 0.001 ***

The strong network activity appeared quickly compared to previously reported human *in vitro* experiments. From hiPSCs to electrophysiological mature co-cultures, our workflow takes about 40 days. Parameters, such as the average mean firing rate and synchrony index, we see at maturity are comparable to human organoids that have been in culture for 30 weeks or cultures that contain rodent cells [10-12]. This is a significant advantage for future studies, as our model is fully human and takes a fraction of the time to reach maturity. Notably, these results were reproducible as seen in Fig 3. We plated the same 2:1 ratio of NPCs to astrocytes on an additional MEA 24-well plate. Recordings reported from plate 2 were 32 DIV, one day less than plate 1. Importantly, plate 2 contained NPCs from two separate differentiation passages compared to plate 1, allowing three different technical replicates of NPCs to be observed. In addition, the two plates contained astrocytes from different passage numbers. Overall, the two plates were similar with parameters such the synchrony index, mean firing rate, and number of active electrodes having no statistical difference. While the network IBI coefficient of variation for plate 1 was about 1.5-fold greater than plate 2, this difference was not statistically significant. However, the average network burst frequency and network burst duration were statistically different between the two plates. These discrepancies could be explained by slight variations in cell numbers due to the low media volumes and cell high density used for plating. Another source of variability could be innate inconsistencies in maturation time or neuronal differentiation. Nevertheless, strong cellular networks formed on both plates in about six weeks, confirming a robust differentiation and maturation protocol.

**Figure 3.**
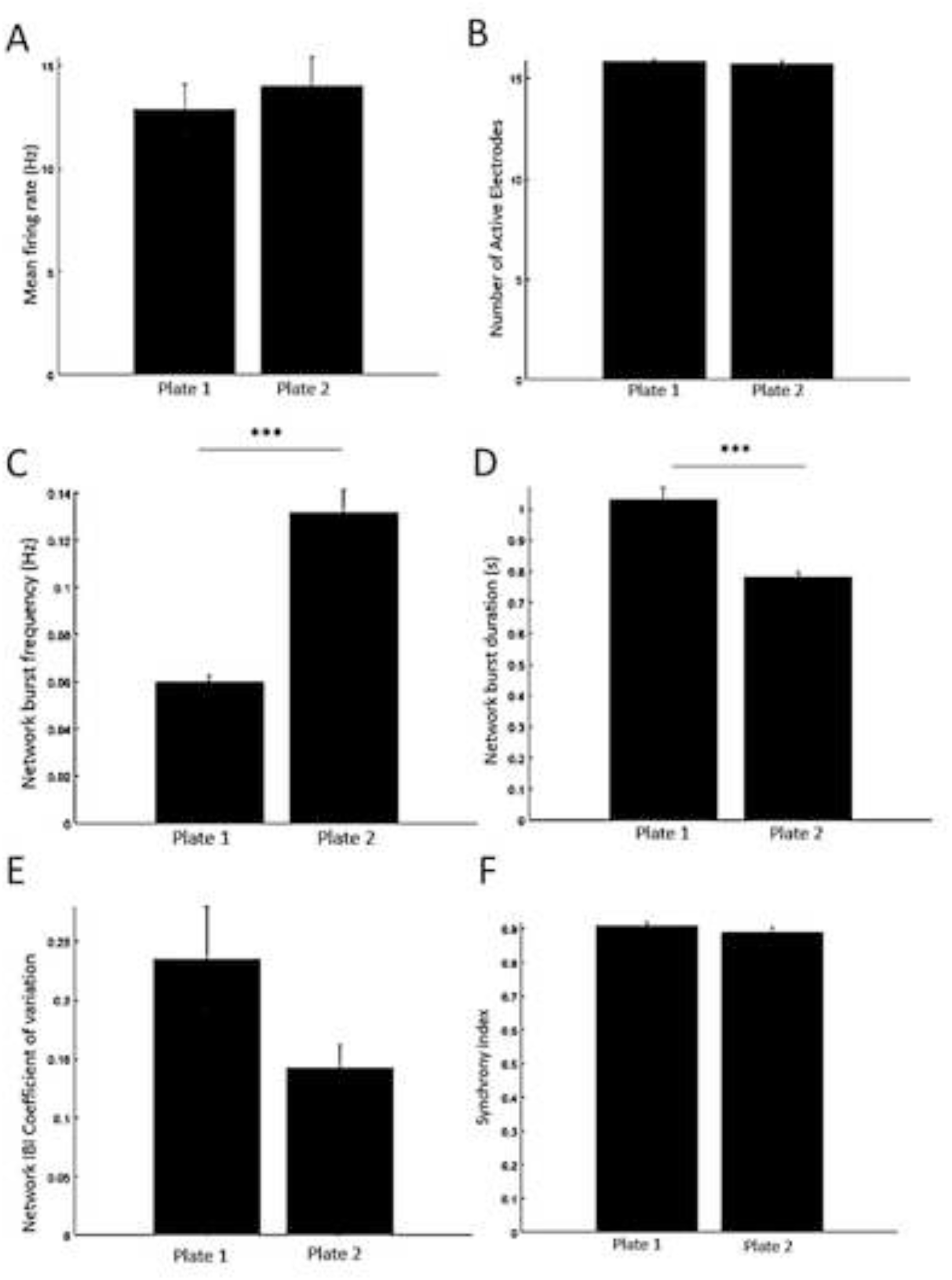
Co-culture electrophysiological reproducibility between two 24-well plates. Characterized by (A) Mean firing rate (Hz) (B) Number of active electrodes (C) Network burst frequency (Hz) (D) Average network burst duration (s) (E) Network interburst interval coefficient of variation (F) Synchrony index. Statistical analysis was performed with an independent t-test. Error bar: SEM. N=24 independent wells per plate. p-value < 0.05 *, p-value < 0.01 **, p-value < 0.001 ***

### Pharmacological validation

Upon achieving network functionality, the co-cultures were validated pharmacologically. All treatments and control wells were recorded 30 minutes and 1 hour after exposure, followed by recordings taken every 2 hours for 24 hours. A washout consisted of a half-media change, and cells were additionally observed and recorded following the wash. Notably, throughout maturation we noted electrophysiological fluctuations due to media changes. Immediately after receiving new media, the mean firing rate and network burst frequency significantly increased. This could also be due to the plate needing time to stabilize in the instrument. In order to avoid these confounding factors, we performed media changes 24 hours before treatments and gave plates 30 minutes of incubation time in the instrument before recordings.

Control wells were either untreated or subjected to 0.1% DMSO to account for treatments dissolved in this solvent. A half-media change was performed on control wells after the 24-hour recording to mimic the washout period in treated wells. We performed pharmacological experiments with glutamatergic receptor inhibitors (Fig 4). CNQX and AP5 were administered to block both AMPA and NMDA receptors, respectively (10 µM each). This treatment decreased the mean firing rate decreasing 4-fold, and network bursts were completely ablated 4 hours after exposure. The activity remained reduced 24 hours after treatment. However, following the washout period, network bursts returned with firing activity in recovery. The synchrony index was also recovering but was still about 1.5-fold less than the baseline. This result confirmed the presence and necessity of glutamatergic neurons in our synchronized cultures.

**Figure 4.**
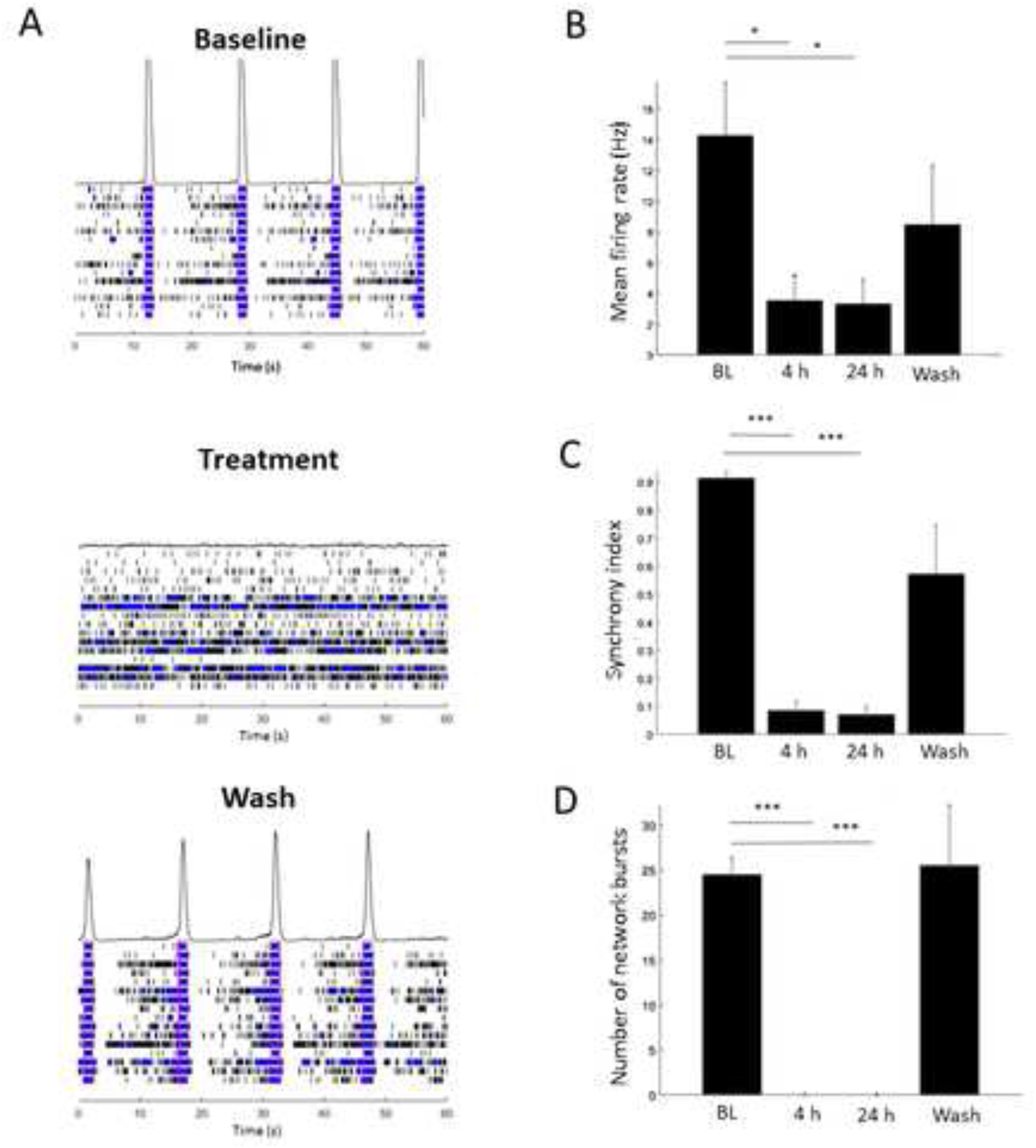
Glutamate receptor inhibitor treatment consisting of 10 µM CNQX and 10 µM AP5. (A) Representative raster plots and histograms of relative population firing rates for baseline (BL), 24 hours after treatment, and after washout. Quantified reductions of (B) Mean firing rate (Hz) (C) Synchrony index (D) Number of network bursts for BL, 4 hours and 24 hours after treatment, and after washout. Statistical analysis was performed with repeated measures ANOVA and Bonferroni corrections. Error bar: SEM. N=4 independent wells. p-value < 0.05 *, p-value < 0.01 **, p-value < 0.001 ***

To test the cell signaling contribution of GABAergic neurons in our cultures, bicuculline (10 µM), a GABA receptor inhibitor, was administered (Fig 5). This produced no significant changes to the cultures at any timepoint, including at the 30-minute incubation not shown. In previous reports using hiPSC cortical derived neurons, 10 µM bicuculline significantly increased the mean firing rate [9]. We did not see this, and in fact had a slight decrease in activity. This result is consistent with the inducible NGN2 protocol producing mainly glutamatergic neurons. In the future, we will perform dose dependent studies using various GABA receptor inhibitors to solidify these findings. This result overall may indicate that the firing patterns we observed were being driven mainly by glutamatergic signaling, and potentially GABAergic neurons present in the culture are not functional in the context of electrophysiological activity. In fact, the absence of inhibitory neuron involvement may explain why we achieved a high mean firing rate so quickly compared to other human *in vitro* models.

**Figure 5.**
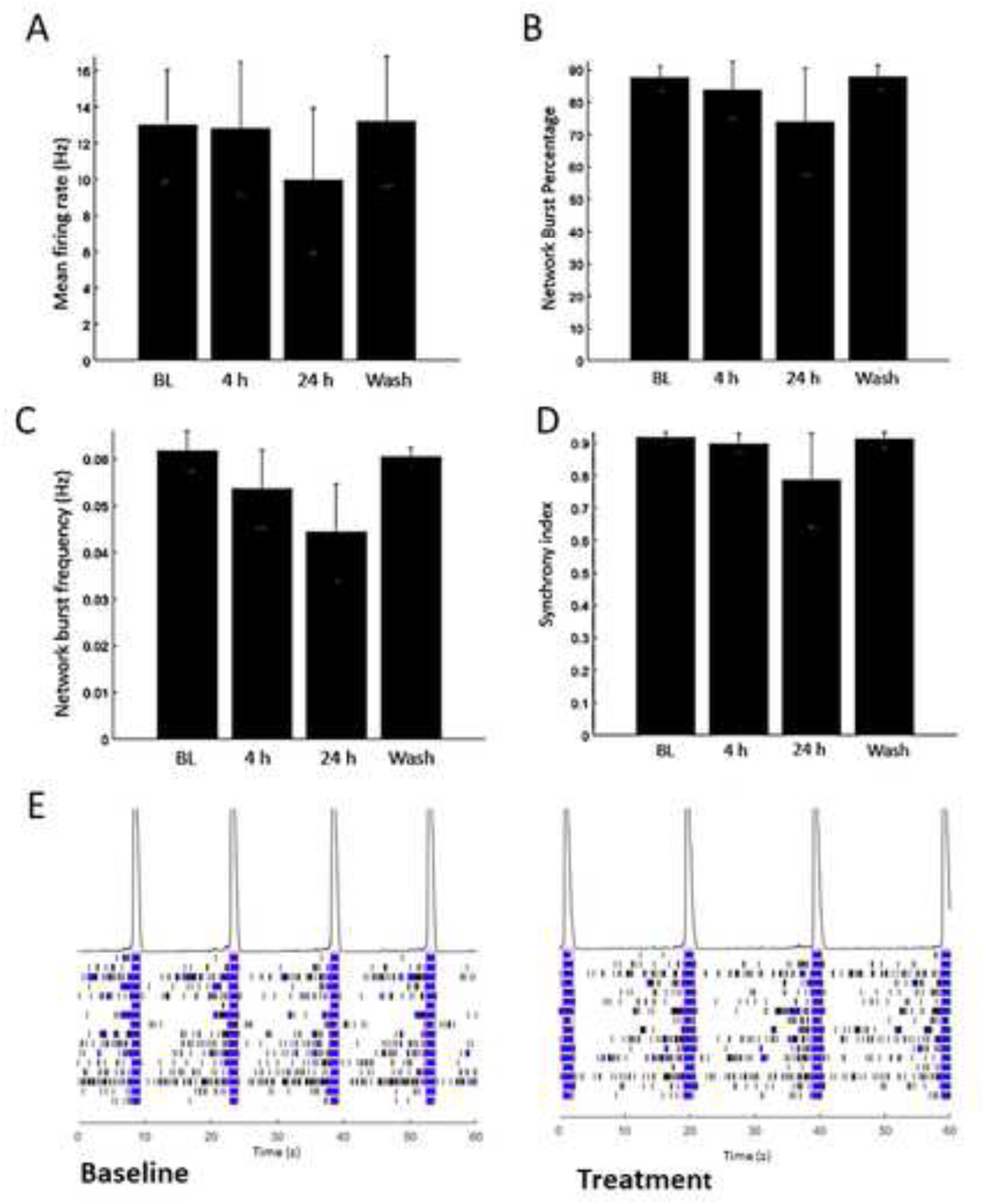
GABA receptor inhibitor treatment consisting of 10 µM bicuculline. Quantifications of (A) Mean firing rate (Hz) (B) Network burst percentage (C) Network burst frequency (D) Synchrony index for baseline (BL), 4 hours and 24 hours after treatment, and after washout. (E) Representative raster plots and histograms of relative population firing rates for baseline and 24 hours after treatment Statistical analysis was performed with repeated measures ANOVA and Bonferroni corrections. Error bar: SEM. N=4 independent wells. p-value < 0.05 *, p-value < 0.01 **, p-value < 0.001 ***

### Modeling Blood Brain Barrier Breakdown

To model *in vitro* the blood brain barrier leakage observed in sporadic Alzheimer’s disease and other pathological conditions, 10% human serum was added directly to cultures as previously described [14]. As shown in Fig 6, this treatment completely disrupted the activity of the culture as the mean firing rate decreased 5.5-fold after 4 hours of exposure. After 24 hours, the culture was nearing zero activity as the mean firing rate dropped to 0.34 Hz. Only 1 of the 4 treated wells had network bursts at the 4-hour timepoint, and after 24 hours there were no network bursting at all. The synchrony index went from a baseline average of 0.88 to 0.01, and, unlike the pharmacological validation treatments, the number of active electrodes was also significantly reduced due to the serum. In addition, the network disruption was captured as early as the 1-hour timepoint indicated by the IBI coefficient of variation significantly increasing to 0.50. Interestingly, after two washout periods, the cultures were able to recover.

**Figure 6.**
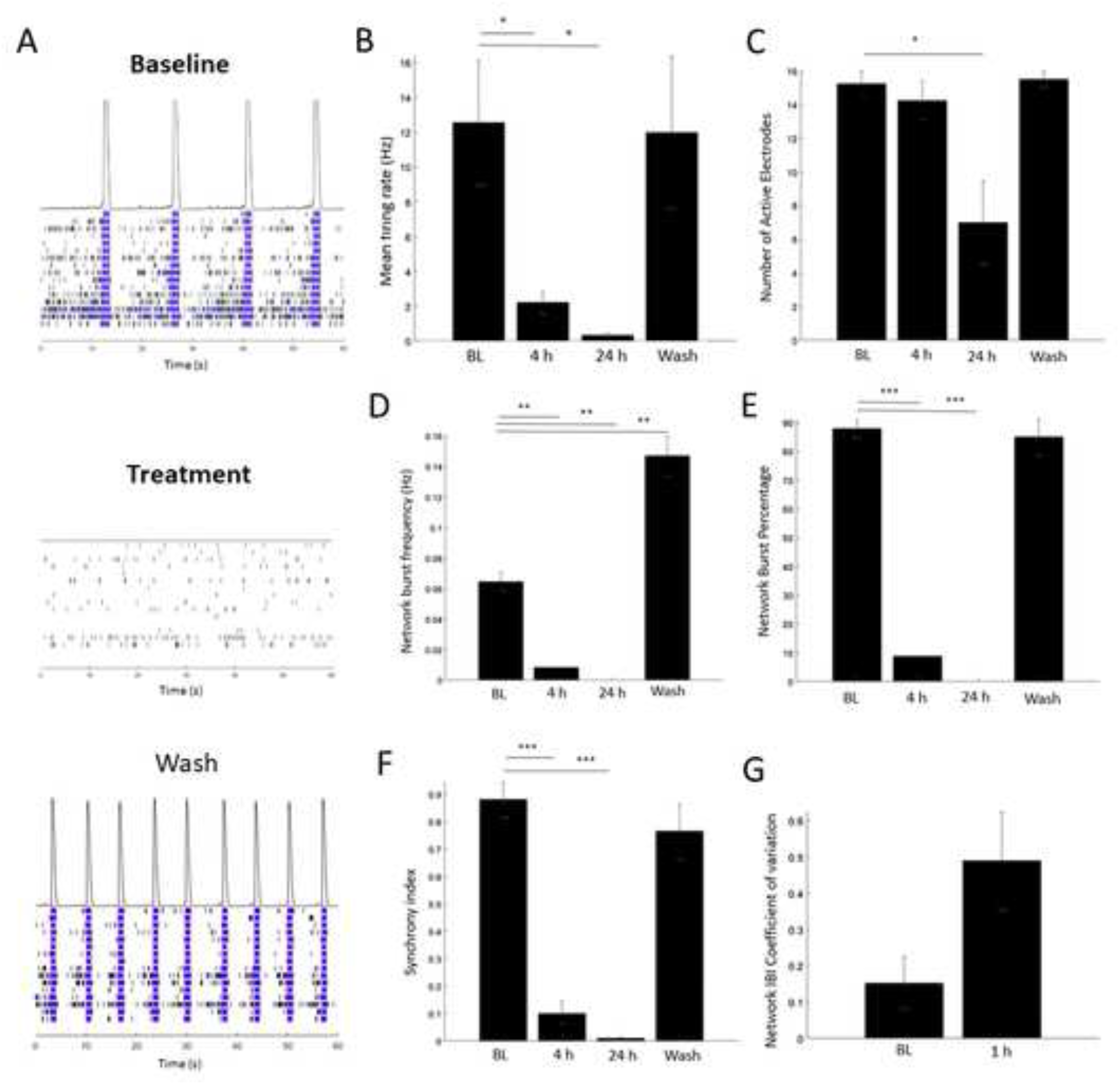
10% human serum treatment. (A) Representative raster plots and histograms of relative population firing rates for baseline, 24 hours after treatment, and after 2 washouts. (B) Mean firing rate (Hz) (C) Number of active electrodes (D) Network burst frequency (Hz) (E) Network burst percentage (%) (F) Synchrony Index (G) Network Interburst Interval Coefficient of Variation. Statistical analysis was performed with repeated measures ANOVA and Bonferroni corrections. Error bar: SEM. N=4 independent wells. p-value < 0.05 *, p-value < 0.01 **, p-value < 0.001 ***

## Discussion

In this study we used neurogenin 2 inducible human pluripotent stem cells differentiated into neurons co-cultured with primary human astrocytes to study electrophysiological parameters associated with the human brain. Differentiated iPSCs were morphologically mature as seen by MAP2 and beta tubulin III staining, and co-cultures were established at a 2 neuron: 1 astrocyte ratio. Spontaneous activity was observed by multielectrode array analysis. We determined that our fully human 2D cultures were able to achieve strong network burst synchrony similar to human organoids in a matter of several weeks compared to previously reported months of maturation.

We confirmed the functionality of our cultures by treating them pharmacologically. Glutamate receptor inhibitors significantly reduced synaptic activity, confirming their role in our network burst synchrony. However, when cultures were treated with a GABA receptor inhibitor, there were no statistically significant changes in any parameter measured in this study. It has been previously reported that iPSCs containing the NGN2 transgene differentiate into largely excitatory neurons (90% glutamatergic) [16, 19, 20]. Therefore, GABAergic neurons present in our culture may not be functional or contributing significantly to the activity of the cultures.

Importantly, we observed electrophysiological reproducibility using our co-culture. These robust results were obtained from three independent differentiations of the NGN2 iPSCs. It has been previously reported that mean firing rate is one of the most variable MEA parameters [21], and our co-culture was consistent not only from well to well but also from plate to plate. In addition, we found using the 2:1 neurons/astrocyte ratio specifically produced the least variability compared to 6:1 and 4:1 (data not shown). Alterations in cell density have been reported to affect parameters like interburst interval [21]. Because of the high-density plating required for this assay, slight variations could be introduced, which could explain why we observed minor variations in some of the parameters comparing plate 1 to plate 2 in Fig 3.

This NGN2 iPSC/primary astrocyte model can also be used to mimic aspects of sporadic Alzheimer’s disease. Human serum exposure has been used to model blood brain barrier leakage reported in AD patients. A previous report has shown 10% serum treatment induces A*β*-like pathology, elevated level of phosphorylated tau, synaptic loss, impaired neural network, and neuronal death after 12 days of exposure [14]. In our studies, network activity was lost after serum exposure. However, cells were able to recover after washout periods with an increased network burst frequency compared to baseline. The specific mechanism by which serum exposure drastically alter electrophysiological parameters is still being explored. Our results could be due to the difference between acute and chronic exposure, and more experiments should be done to determine what compound(s) in the serum itself are causing these drastic electrophysiological changes. One possible serum component is β2-microglobulin. A recent publication found elevated levels of this protein in human down syndrome plasma. When β2-microglobulin was administered to wild type mice, they observed synaptic and memory deficits citing NMDA receptor dysfunction [22].

One additional interesting feature of these co-cultures is functional network connectivity that has been referred to as “sleep in a dish”. Synchronous network burst frequency of cell culture models may resemble aspects of slow wave sleep *in vivo* [23-24]. Specifically, “sleep in a dish” is described as a low frequency burst-pause-burst pattern in the range of 0.06-0.2 Hz [23, 25-26]. This pattern is the default state of isolated cells that has been well established in dissociated rodent cortical primary cultures and more recently in iPSCs [25-26]. Our findings replicate these earlier studies and suggest this model may be a useful *in vitro* approach to explore the bidirectional relationship between Alzheimer’s disease and sleep disturbances.

In conclusion, this fully human co-culture exhibits strong, synchronous network bursting on multielectrode arrays in less time than previously reported. We were able to investigate aspects of human brain network function and model characteristics of Alzheimer’s disease. Using MEA electrophysiology is beneficial as it can sensitively measure neuronal dysfunction, not just assay neuronal death. This technique can also mimic aspects of clinical data without harming human health. Therefore, this method provides a relatively cost effective, fast, high throughput, human method for pharmacological testing of potential treatments for neurodegenerative diseases.

## Acknowledgements

This work was performed in the Stem Cell Research and Technology Resource Center at the University of Colorado Boulder. We would like to thank Dr. Roy Parker and his lab for advising us on the hiPSC differentiation protocol. This project was funded by National Institute on Aging of the National Institutes of Health award RF1AG064465 to MRO, CH, and CDL.

## Notes

### Competing Interest Statement

The authors have declared no competing interest.

